# Odor modality is transmitted to cortical brain regions from the olfactory bulb

**DOI:** 10.1101/2023.02.26.530150

**Authors:** Michelle F. Craft, Andrea K. Barreiro, Shree Hari Gautam, Woodrow L. Shew, Cheng Ly

## Abstract

Odor perception is the impetus for important animal behaviors with two predominate modes of processing: odors pass through the front of the nose (orthonasal) while inhaling and sniffing, or through the rear (retronasal) during exhalation and while eating. Despite the importance of olfaction for an animal’s well-being and that ortho and retro naturally occur, it is unknown how the modality (ortho versus retro) is even transmitted to cortical brain regions, which could significantly affect how odors are processed and perceived. Using multi-electrode array recordings in tracheotomized anesthetized rats, which decouples ortho-retro modality from breathing, we show that mitral cells in rat olfactory bulb can reliably and directly transmit ortho versus retronasal modality with ethyl butyrate, a common food odor. Drug manipulations affecting synaptic inhibition via GABA_A_ lead to worse decoding of ortho versus retro, independent of whether overall inhibition increases or decreases, suggesting that the olfactory bulb circuit may naturally favor encoding this important aspect of odors. Detailed data analysis paired with a firing rate model that captures population trends in spiking statistics shows how this circuit can encode odor modality. We have not only demonstrated that ortho/retro information is encoded to downstream brain regions, but also use modeling to demonstrate a plausible mechanism for this encoding: due to synaptic adaptation, it is the slower time course of the retronasal stimulation that causes retronasal responses to be stronger and less sensitive to inhibitory drug manipulations than orthonasal responses.

**New and Noteworthy:** Whether ortho (sniffing odors) versus retro (exhalation and eating) is encoded from the olfactory bulb to other brain areas is not completely known. Using multi-electrode array recordings in anesthetized rats, we show that the olfactory bulb transmits this information downstream via spikes. Altering inhibition degrades ortho/retro information on average. We use theory and computation to explain our results, which should have implications on cortical processing considering that only food odors occur retronasally.

## Introduction

Olfaction is driven by odor molecules entering the nasal cavity, inducing a cascade of action potentials in the nervous system that transmit and process odor information. There are two routes by which odor molecules can enter the nasal cavity: through the nostrils during inhalation and sniffing (**ortho)**, or from the throat during exhalation and while eating **(retro**). Orthonasal stimulation is by far the most commonly studied modality, despite the fact that retonasal delivery naturally occurs during eating. Prior imaging studies have shown differences in the activation of the regions of the nasal epithelium with ortho and retro stimulation (Sanganahalli et al., 2020; Gautam and Verhagen, 2012b; Furudono et al., 2013). Moreover, humans are able to discriminate whether food odors are delivered ortho or retronasally without being told which modality and without active sniffing (Frasnelli et al., 2008). Imaging studies have shown that at least with food odors, ortho/retro are two distinct modalities, rather than 2 different routes to the same modality, independent of odor intensity (Bender et al., 2009). Importantly, there is evidence that cortical processing depends on whether odors are delivered ortho- versus retro-nasally (Blankenship et al., 2019; Gautam and Verhagen, 2012a), and humans report perceiving the same smells differently depending on ortho or retro modality (Hannum et al., 2021; Small et al., 2005). Together these studies convincingly show that ortho versus retro information is transmitted to cortical brain regions, but the details of how this occurs and the core network mechanisms that facilitate this encoding are largely unknown.

There are several ways that ortho/retro information could be made available to higher brain regions. Since ortho and retro stimuli are naturally dictated by inhalation and exhalation, the motor signals that control breathing could provide efferent copies of this information to various brain regions. Moreover, even without such motor signals, the mechanical stimulation of neurons in the nasal cavity by breath-driven airflow generates strong breath related signals throughout the olfactory system (Iwata et al., 2017; Shusterman et al., 2011). However, previous experiments have delivered stimuli independently of the breath cycle, and still found significant differences in ortho versus retro response in cortex (Small et al., 2005; Heilmann and Hummel, 2004) and ORN input to OB (Gautam and Verhagen, 2012b). Thus, there are important breath-independent aspects of ortho/retro coding. Although these different mechanisms for odor modality encoding are all plausible, here we designed our experiments to study the ‘bottom up’, breath-independent mechanisms. We specifically performed a double tracheotomy which redirected the rats’ breath to bypass the nasal cavity entirely and allowed us to deliver ortho and retro stimuli at times that were independent of inhale and exhale times. So the mechanisms of ortho and retro coding that we study here are independent of any motor or mechanosensory signals caused by breathing.

Before odor information reaches the brain, it is processed in the olfactory bulb (**OB)** and relayed to higher brain regions via excitatory mitral cells (**MC**s) (and tufted cells). Thus, the OB is critical for determining whether ortho versus retronasal odors are encoded before being conveyed to the brain for processing and perception. More broadly, OB activity is tied to odor perception (Mandairon and Linster, 2009). While the role of inhibitory cells in influencing OB processing of ortho versus retro odors is unknown, inhibition is known to play a key role in many other aspects of OB processing. OB inhibition is known to alter activity patterns that represent odors (Wilson and Laurent, 2005), granule cells that provide inhibition reflect changes in odor concentration (Tan et al., 2010), and OB inhibition levels alter odor discrimination dynamics (Abraham et al., 2010). These facts motivated us to both decrease and increase the levels of GABA_A_ synaptic inhibition via drug manipulations in our experiments, which consist of *in vivo* recordings of MCs in the OB on double tracheostomized rats.

We show that the mode of olfaction (ortho versus retro) is indeed transmitted to cortical brain regions with a common food odor, ethyl butyrate (**EB)**. We find that encoding is generally good and well above chance level for the intact circuit and even with altered levels of inhibition. Thus downstream brain regions may readily have access to ortho versus retronasal odor modality encoded via firing rate from the OB. Importantly, the altered circuits with both increases (via **Mus**cimol application, a GABA_A_ agonist) and decreases in inhibition (via **Bic**uculline, a GABA_A_ antagonist) result in worse individual encoding on average. This suggests that the intact OB circuit may be advantageous for encoding ortho versus retro modality.

Our data enables investigation of the OB circuit mechanisms that promote this encoding with a computational model. We show how a simple excitatory-inhibitory **(E-I**) reciprocally coupled model captures the modality dependent differences in the population firing rate. Our data shows that the GABA_A_ (ant-)agonists do not significantly alter the odor-evoked population firing rate with slower longer-lasting retronasal stimulation, in contrast to ortho stimulation. Our model captures these aspects of our data, exploiting the temporal differences, i.e., the slow retro stimulation leads to short-term synaptic depression of the excitatory synapse to diminish the effects of the inhibition. Furthermore, we model trial-to-trial spike rate variability to be largely consistent with our data, so that the model can encode orth-/retro-like stimuli. We find that average encoding is degraded when inhibition is either increased or decreased, as observed in our data. Given the commonality of reciprocally coupled excitatory-inhibitory circuits, these results may apply outside of the OB, specifically to where stimuli have temporal differences.

## Materials and methods

### Ethics statement

All procedures were carried out in accordance with the recommendations in the Guide for the Care and Use of Laboratory Animals of the National Institutes of Health and approved by University of Arkansas Institutional Animal Care and Use Committee (protocol #14049). Isoflurane and urethane anesthesia were used and urethane overdose was used for euthanasia.

### Data and code availability

See https://github.com/michellecraft64/Modality for MATLAB code implementing the data analysis and firing rate model.

See https://www.dx.doi.org/10.6084/m9.figshare.14877780 for raw data collected from the Shew Lab.

### Electrophysiological recordings

Data were collected from 8 adult male rats (240-427 g; *Rattus Norvegicus*, Sprague-Dawley outbred, Harlan Laboratories, TX, USA) housed in an environment of controlled humidity (60%) and temperature (23°C) with 12h light-dark cycles. The experiments were performed in the light phase.

#### Surgical preparations

Anesthesia was induced with isoflurane inhalation and maintained with urethane (1.5 g/kg body weight (**bw)** dissolved in saline, intraperitoneal injection (**ip)**). Dexamethasone (2 mg/kg bw, ip) and atropine sulphate (0.4 mg/kg bw, ip) were administered before performing surgical procedures. Throughout surgery and electrophysiological recordings, core body temperature was maintained at 37°C with a thermostatically controlled heating pad. To isolate the effects of olfactory stimulation from breath-related effects, we performed a double tracheotomy surgery as described previously (Gautam and Verhagen, 2012b). A Teflon tube (OD 2.1 mm, upper tracheotomy tube) was inserted 10 mm into the nasopharynx through the rostral end of the tracheal cut. Another Teflon tube (OD 2.3 mm, lower tracheotomy tube) was inserted into the caudal end of the tracheal cut to allow breathing, with the breath bypassing the nasal cavity. Both tubes were fixed and sealed to the tissues using surgical thread. Local anesthetic (2% Lidocaine) was applied at all pressure points and incisions. Subsequently, a craniotomy was performed on the dorsal surface of the skull over the right olfactory bulb (2 mm × 2 mm, centered 8.5 mm rostral to bregma and 1.5 mm lateral from midline).

#### Olfactory stimulation

A Teflon tube was inserted into the right nostril to deliver orthonasal stimuli, and the left nostril was sealed by suturing. The upper tracheotomy tube inserted into the nasopharynx was used to deliver odor stimuli retronasally. Odorized air was delivered for 1 second in duration at 1 minute intervals, with a flow rate of 250 ml/min and 1% of saturated vapor. Two odors were used: ethyl butyrate **(EB**, a food odor) and 1-Hexanol (**Hex**, a non-food odor). Here we limited our analysis to EB trials because food odors are perceived ortho- and retro-nasally (Small et al., 2005); non-food odors do not naturally occur retronasally.

#### Electrophysiology

A 32-channel microelectrode array (MEA, A4x2tet, NeuroNexus, MI, USA) was inserted 400 *µ*m deep from dorsal surface of OB targeting tufted and mitral cell populations. The MEA probe consisted of 4 shanks (diameter: 15 *µ*m, inter-shank spacing: 200 *µ*m), each with eight iridium recording sites arranged in two tetrode groups near the shank tip (inter-tetrode spacing: 150 *µ*m, within tetrode spacing 25 *µ*m). Voltage was measured with respect to an AgCl ground pellet placed in the saline-soaked gel foams covering the exposed brain surface around the inserted MEAs. Voltages were digitized with 30 kHz sample rate (Cereplex + Cerebus, Blackrock Microsystems, UT, USA). Recordings were band-pass filtered between 300 and 3000 Hz and semiautomatic spike sorting was performed using Klustakwik software, which is well suited to the type of electrode arrays used here (Rossant et al., 2016).

#### Pharmacology

We topically applied GABA antagonists (bicuculline 20 *µ***M**) or agonists (muscimol 20 *µ*M) dissolved in artificial cerebrospinal fluid (**ACSF**). Gel foam pieces soaked in the ACSF+drug solutions were placed on the surface of OB surrounding the recording electrodes. Note that the concentration of (ant-)agonists can vary over orders of magnitude; some labs use 10 *µ*M (Li et al., 2019; Sharp and Finger, 2002) and others use up to 10 to 10^4^ higher concentration levels (Wachowiak and Cohen, 1999; Kollo et al., 2014). The gel foams were placed 10 minutes before beginning a recording.

### Data analysis

Data was collected *in vivo* from the mitral cell (**MC**) layer in the olfactory bulb (**OB**) of multiple anesthetised rats using a multi-electrode array recording. The data consisted of spike recordings of multiple MC spike responses to EB and Hex presented by the two routes of stimulation, orthonasally and retronasally, for a total of 20 trials, 10 for each ortho and retro, with the order alternating (10 ortho before 10 retro, then 10 retro before 10 ortho trials). The spike counts were calculated using 100 ms overlapping time windows. Three separate drug preparations were used in order to analyze inhibitory effects on MC spiking responses: no drug (control), Bicuculline (GABA_A_ antagonist, i.e., decreasing inhibition), and Muscimol (GABA_A_ agonist, i.e., increasing inhibition). Data was obtained from 8 viable rats in total. After spike sorting, we identified and eliminated units (cells) that had firing rates below 0.008 Hz or more than 49 Hz, calculated over the <u>entire recording duration</u>. We took this conservative approach to eliminate potential artifacts yielding units that had unrealistically high or low firing; see Table 2 for the reported number of total rats and individual cells for each drug preparation. Note that some cells around the odor onset (*t* = 0) will have much larger (*>* 49 Hz) or smaller firing rates than our criteria to eliminate units, but the <u>entire recording duration</u> is much longer with approximately a minute between trials.

### Defining single cell decoding

We used a form of linear discriminant analysis (**LDA**) for decoding accuracy. Decoding accuracy is defined as the fraction of trials correctly classified by a threshold that maximizes decoding accuracy. Letting *s*_*k*_(*t*) be the spike train of the *k*^*th*^ trial (*k* = 1, 2, …, *N*) that at each time *t* is either = 0 (no spike) or = *δ*(*t* − *t*_*j,k*_), where *t*_*j,k*_ denotes the *j*^*th*^ spike on the *k*^*th*^ trial . The entries of the spiking rate 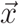 are:

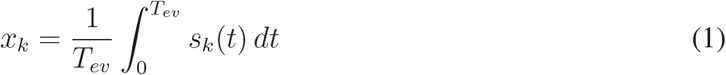

where *t* = 0 is the time the odor is presented. There are *N* = 20 trials total (10 ortho, 10 retro) for each individual cell. We consider *N* different threshold values *θ*_*k*_:

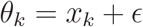

and use both classification schemes (decode stimulus as ortho if *x*_*k*_ *< θ* and retro if *x*_*k*_ *> θ*, or as ortho if *x*_*k*_ *> θ*). From this comprehensive list, we select an optimal threshold *θ*^∗^ and classification scheme that results in the best decoding accuracy. Here *ϵ* ≈ 10^−16^ is machine precision and is there to insure that each trial is categorized as ortho or retro. Since decoding accuracy is the percent of correctly classified trials (ortho or retro) and *N* = 20, it varies between 0.5 (chance) and up to 1 in increments of 0.05 for each cell.

To systematically compare differences in decoding accuracy, we select a fixed time window for all drug preparations and modalities based on two statistical tests (two sample *t*−test assuming unequal variances and Wilcoxon rank sum test) and their *p*−values. We evaluated decoding accuracy from *T*_*ev*_ =100 ms to *T*_*ev*_ =1 s in 100 ms increments in the evoked state for each drug preparation. We analyzed differences in mean between no drug preparation and Bicuculline, as well as no drug preparation and Muscimol. This resulted in 4 different *p*−values for each time window; we found that *T*_*ev*_ = 900 ms had the most significant *p*−values for both combinations of no drug/Bicuculline and no drug/Muscimol, see Fig A2A,C. Figure A2B,D shows the population decoding accuracies (mean across the population and one standard deviation in shaded region).

### Statistical significance

We use three tests to assess whether a given effect is statistical significant: 1) two-sample *t*−test with unequal variance, 2) Wilcoxon rank sum test, 3) one-way ANOVA. Each of these tests are used to rule out the null hypothesis that the population means are the same in two different categories (Cohen, 2013). Each test is different and has various underlying assumptions: 1) normal samples, 2) non-parametric test assuming independent groups and equal variance, 3) including sample variance to assess differences in means assuming equal variance and normally distributed residuals. We consider three tests to provide a more complete picture of the results and to demonstrate robustness, or lack thereof, for a given effect.

In addition to the *p*−values that indicate the probability of the null hypothesis holding (no effect, same population means), we also report the effect sizes. We use the conventions outlined in Cohen (2013); Tomczak and Tomczak (2014) that provide formulas for effect sizes, and a qualitative ‘rule of thumb’ for when effect sizes are: **small, medium** or **large**. For 2 groups of length *n*_1_ and *n*_2_ respectively, the effect sizes we use:

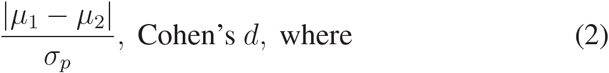

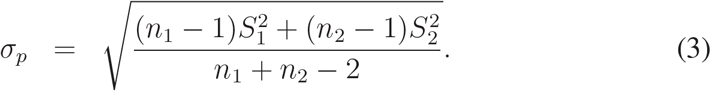

For Wilcoxon rank sum test, use:,

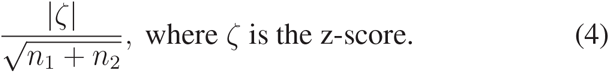

For one-way ANOVA, use:

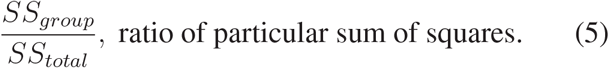

The variables *µ*_*j*_ and 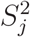 denote the sample mean and variances of the *j*^*th*^ group. Effect sizes for *t*−test and Wilcoxon rank sum test with values: (0, 0.2] are considered **small**, (0.2, 0.5] are **medium**, (0.5, 0.8] and above are **large** (Cohen, 2013; Tomczak and Tomczak, 2014). Effect sizes for one-way ANOVA with values (0, 0.01] are considered **small**, (0.01, 0.06] are **medium**, (0.06, 0.14] and above are l**arge** (Cohen, 2013).

### Population decoding

The two methods are applied to each population of simultaneously recorded MCs.

PCA+LDA: For each recording, Principle Components Analysis (**PCA**) was applied to the concatenated matrix *X* of size 20 × *M*, where *M* is the number of MCs (Fig 6A), rows 1–10 correspond to the spiking rate with ortho and rows 11–20 correspond to retro. We only use the first two principal components and apply Linear Discriminate Analysis **(LDA)** to find a line in the 2-dimensional plane that maximizes classification accuracy (Cunningham and Yu, 2014). We used built-in MATLAB routines `pca, fitcdiscr, predict` (see GitHub for code).

SVM: We use support vector machine with supervised ‘learning’ to classify data into ortho or retro. We use a nonlinear classification method with Gaussian kernels, and Bayesian optimization to find the best kernel scale and box constraints for the SVM (we insured that for all recordings and drug preparations, the objective was minimized within the allowable function evaluations). After the SVM is fit to the data, the decoding accuracy is the correct classification rate obtained from the MATLAB function `kfoldLoss` using 10-fold cross-validation. We used built-in MATLAB routines (see GitHub link for code).

### Firing rate model

We use a space-clamped Wilson-Cowan rate model of coupled E-I cells: E (MC, excitatory) and I (inhibitory from PGC, GC, etc). The models of the respective cell firing rate, *A*_*j*_(*t*) *j* ∈ (*M, P*), are represented by the following ordinary differential equations:

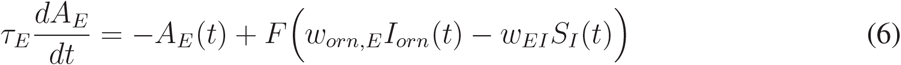

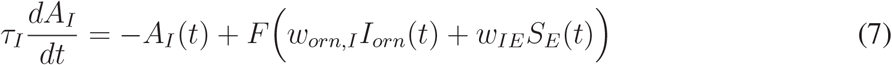

where the synaptic term, *S*_*j*_ *j* ∈ {*E, I*}, is defined by synapses with rise and decay time scales (same *τ*_rise_, *τ*_decay_ for E/I) as follows:

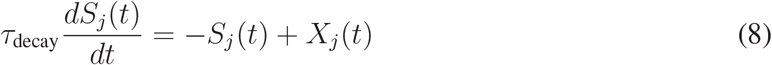

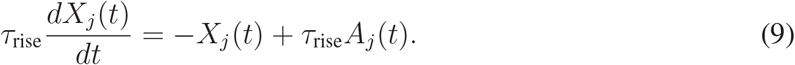

In Eqs 6 and 7, *I*_*j*_(*t*) is the sum of external currents (e.g., stimulus input and other currents) that varies over time to account for stimulus input, *w*_*orn,j*_ are the coupling strengths of olfactory receptor neuron (**ORN**) input to *j, w*_*jk*_ are the coupling strengths from cell *k* to *j*, and the transfer function *F* is a threshold linear function:

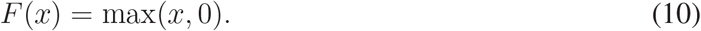

Without synaptic depression, we fix *w*_*IE*_ = 1. With synaptic depression, *w*_*IE*_(*t*) is governed by:

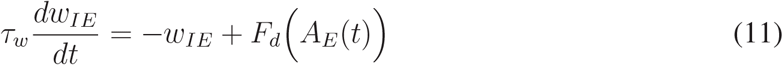

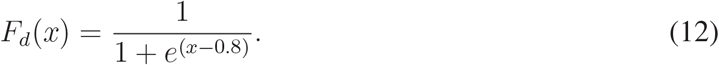

with initial value *w*_*IE*_(0) set to its steady-state value at *t* = −0.5 s in Eqs(6)–(12).

The trial-to-trial variability of spiking rate is modeled as a negative binomial random variable:

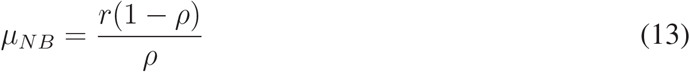

Given output from the Wilson-Cowan model *A*_*E*_(*t*), we set:

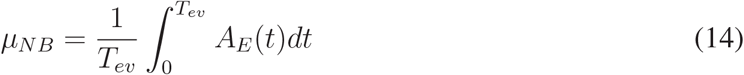

by manually determining the *ρ* parameter in the negative binomial distribution (see Table 1) for

**Table 1:** Parameter values in firing rate model (Eqs (6)–(12)). Last 2 rows describe the parameter *ρ* in simulated trial variability using negative binomial distribution.

| Parameter | Value |  |  |
| --- | --- | --- | --- |
| $\tau_E$ | 10 ms | | |
| $\tau_I$ | 5.5 ms | | |
| $\tau_{decay}$ | 10 ms | | |
| $\tau_{rise}$ | 2 ms | | |
| $\tau_w$ | 100 ms | | |
| $w_{orn,E}$ | 1 | | |
| $w_{orn,I}$ | 1.1875 (ortho), 0.3 (retro) | | |
|  | Drug Preparation: |  |  |
|  | No Drug | Bic | Mus |
| $w_{EI}$ | 0.2 | 0.15 | 0.25 |
| $\rho$ for ortho | 0.32 | 0.32 | 0.32 |
| $\rho$ for retro | 0.32 | 0.53 | 0.052 |

**Table 2:** Number of rats and respective individual cells for each drug preparation subject to the food odor ethyl butyrate (EB).

|  | No Drug | Bicuculline | Muscimol |
| --- | --- | --- | --- |
| Rats | 8 | 4 | 3 |
| Cells | 913 | 413 | 419 |

each drug preparation, and setting the parameter *r* = *µ*_*NB*_*ρ/*(1 − *ρ*).

## Results

We collected spike data with multi-electrode array recordings of urethane anesthetized rats from the OB mitral cell layer, where each cell (MC) was subject to the same odor delivered with ortho and retro stimulation (Fig 1Ai), repeated for 10 trials for each modality. In addition, two drugs were applied to alter the circuit: a GABA_A_ antagonist (**Bic**uculline) and a GABA_A_ agonist (**Mus**cimol), see Table 2. For reference, Fig 1Aii shows the population-averaged firing rate (trial-average and population-averaged firing rate) by modality for a given drug preparation, with *t* = 0 denoting time of odor presentation, held for 1 s. This is commonly referred to as the peri-stimulus time histogram (**PSTH**).

**Figure 1:**
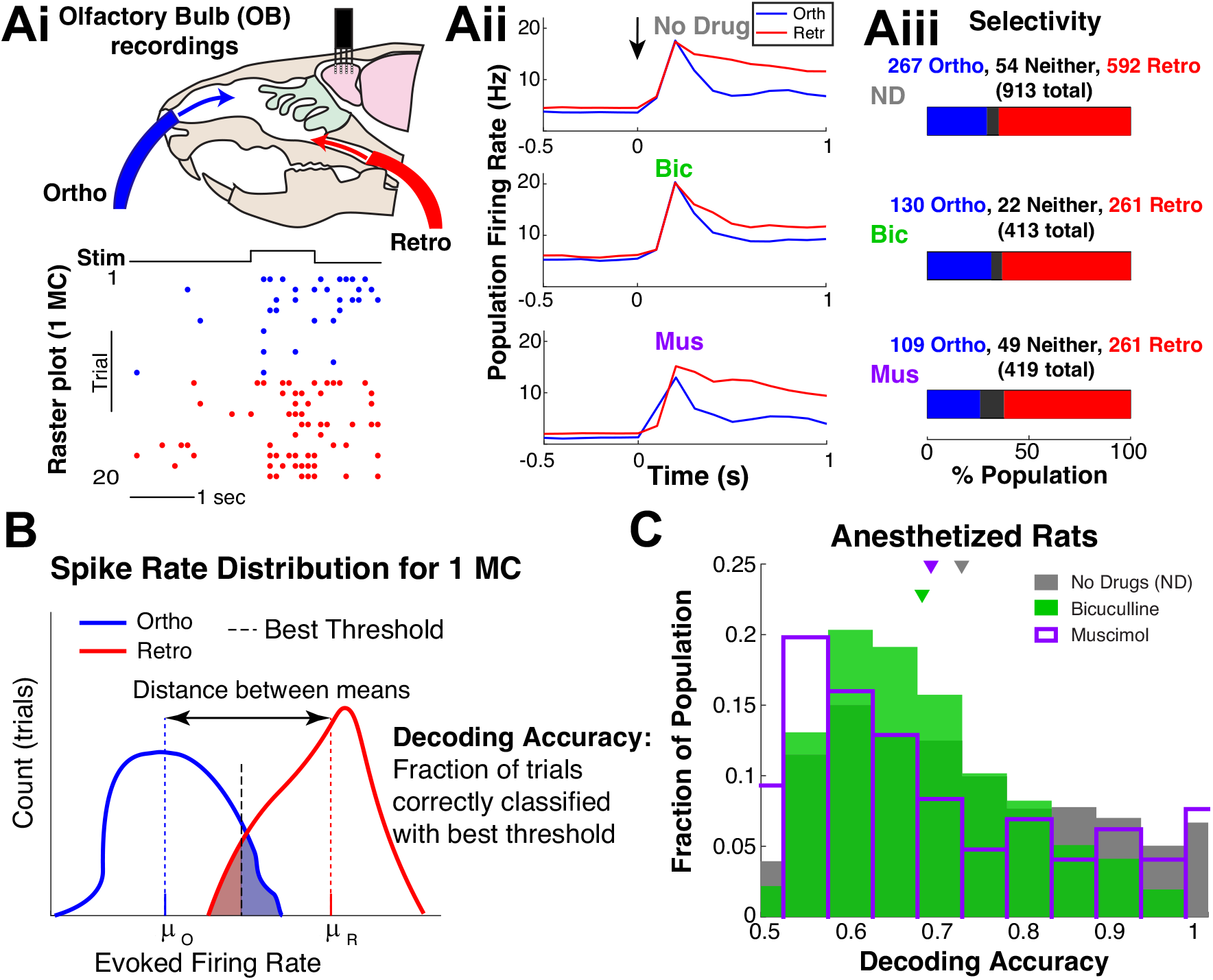
Individual mitral cells encode modality. Ai) Experiment setup to test whether OB MCs encode ortho versus retro stimulation, with example raster plot for 1 MC. Aii) Population-averaged firing rate (trial- and population-averaged), i.e., **PSTH** for each modality and drug preparation; *t* = 0 denotes when ethyl butyrate (food odor) is presented. **Aiii**) The proportion of population that fire more for ortho (or retro), i.e., ortho selective ⇔ higher trial averaged firing rate for ortho than retro. **B**) Schematic of how decoding accuracies are calculated for each MC based on trial firing rates: *µ*_*R/O*_ are the trial-averaged firing rates. Shaded regions correspond to incorrect decoding, unshaded regions to correct decoding. C) Distribution of decoding accuracies of MC with 3 different preparations: intact no drug (gray), less inhibition via Bic (green) and more inhibition via Mus (purple). Mean decoding accuracy for no drug is 0.724; with Bic and Mus the means are: 0.680, 0.690 (resp.). Differences are all statistically significant (*α* = 0.01 with twosample *t*−test assuming unequal var, Wilcoxon rank sum test, and one-way ANOVA) with mostly ‘medium’ effect size, see Table 3. Windows were determine systematically to maximize *p*−values in testing average decoding accuracy, see Fig A2A,C and **Materials and methods.**

### Individual MCs encode ortho versus retro

To remain agnostic to how higher brain regions decode population activity, we predominately consider individual MCs odor modality encoding, a logical first step (Rolls and Treves, 2011). We use odor-evoked firing rate in a given trial as the source of encoding. Specifically, the firing rate in the *k*^*th*^ trial, *x*_*k*_, is the sum of the evoked spike counts normalized by time window *T*_*ev*_ = 0.9 s:

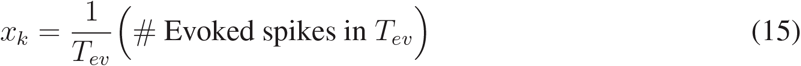

for *k* = 1, 2, …, 20, same as Eq (1) in **Materials and methods**. The time window *T*_*ev*_ is systematically chosen from a range of possible values, see Fig A2A,C in Appendix and **Materials and methods** for details. To assess whether MCs might prefer one modality over the other, we compare the trial-averaged firing rate for ortho and retro as a measure of selectivity (Fig 1Aiii). Most MCs ‘prefer’ or spike more with retro than ortho (≈ 63 %), and the proportions do not vary much across drug preparations (but see Appendix and Table A1 for analysis of Off-Response cells).

We define decoding accuracy as the fraction of trials correctly classified by a threshold that maximizes decoding accuracy; this is a measure of information accessible by an ideal observer from each MC individually (Fig 1B). We only consider EB, a common food odor, because typically only food odors are perceived retronasally (Small et al., 2005). It has been shown that nonfood odors artificially delivered retronasally can be perceived as well as food odors (Small et al., 2005). However, numerous studies have shown that human perception of non-food odors delivered retronasally is degraded compared to food odors (Hannum et al., 2021; Bender et al., 2009; Frasnelli et al., 2008).

The OB circuit encodes ortho versus retro very well. Figure 1C shows the spiking activity of individual MCs encode (measured by decoding accuracy) ortho versus retro for the vast majority of cells. MCs have a wide range of decoding accuracies varying between 0.5 (chance) and 1 (perfect decoding). With no drugs, the mean decoding accuracy (0.724) is better than both Bic and Mus, which have lower average decoding accuracies of 0.680, 0.690, respectively. These averages are numerically small in difference, but note that the range of decoding accuracies is only 0.5. Furthermore, the differences in the mean decoding accuracy are all statistically significant (*α* = 0.01) using three tests: two-sample *t*−test, Wilcoxon rank sum test, and one-way ANOVA. Table 3 demonstrates that the *p*−values are all small, and that the effect size is generally ‘medium’, with some ‘small’ (see Statistical significance section in **Materials and methods)**.

**Table 3:** Population Decoding Accuracy: significance measured by *p*−values of average decoding accuracies over MC, shown in Figure 1C. Using 3 tests: two-sample *t*−test assuming unequal variance, Wilcoxon rank sum test, and one-way ANOVA. Bottom table shows effect size (see **Materials and methods**).

| Relationship / $p$-value | $t$-test | Rank Sum | ANOVA |
| --- | --- | --- | --- |
| Bic < ND | $3.6 \times 10^{-10}$ | $4.9 \times 10^{-6}$ | $1.8 \times 10^{-8}$ |
| Mus < ND | $1.2 \times 10^{-4}$ | $1.8 \times 10^{-6}$ | $7.1 \times 10^{-5}$ |

| Relationship / Effect Size | $t$-test | Rank Sum | ANOVA |
| --- | --- | --- | --- |
| Bic < ND | 0.336 (medium) | 0.126 (small) | 0.0237 (medium) |
| Mus < ND | 0.235 (medium) | 0.131 (small) | 0.0118 (medium) |

Figure 2 shows a more complete picture of the entire (pooled) population’s spike rate statistics across drug preparations and modality, beyond the PSTH in Figure 1Aii. There are very strong positive correlations between the mean firing rates and standard deviations (across trials) in all drug preparations and both modalities (see Figure A3, and Table A2). For further discussion on the modulation of evoked spiking variability in the OB, see Ly et al. (2021).

**Figure 2:**
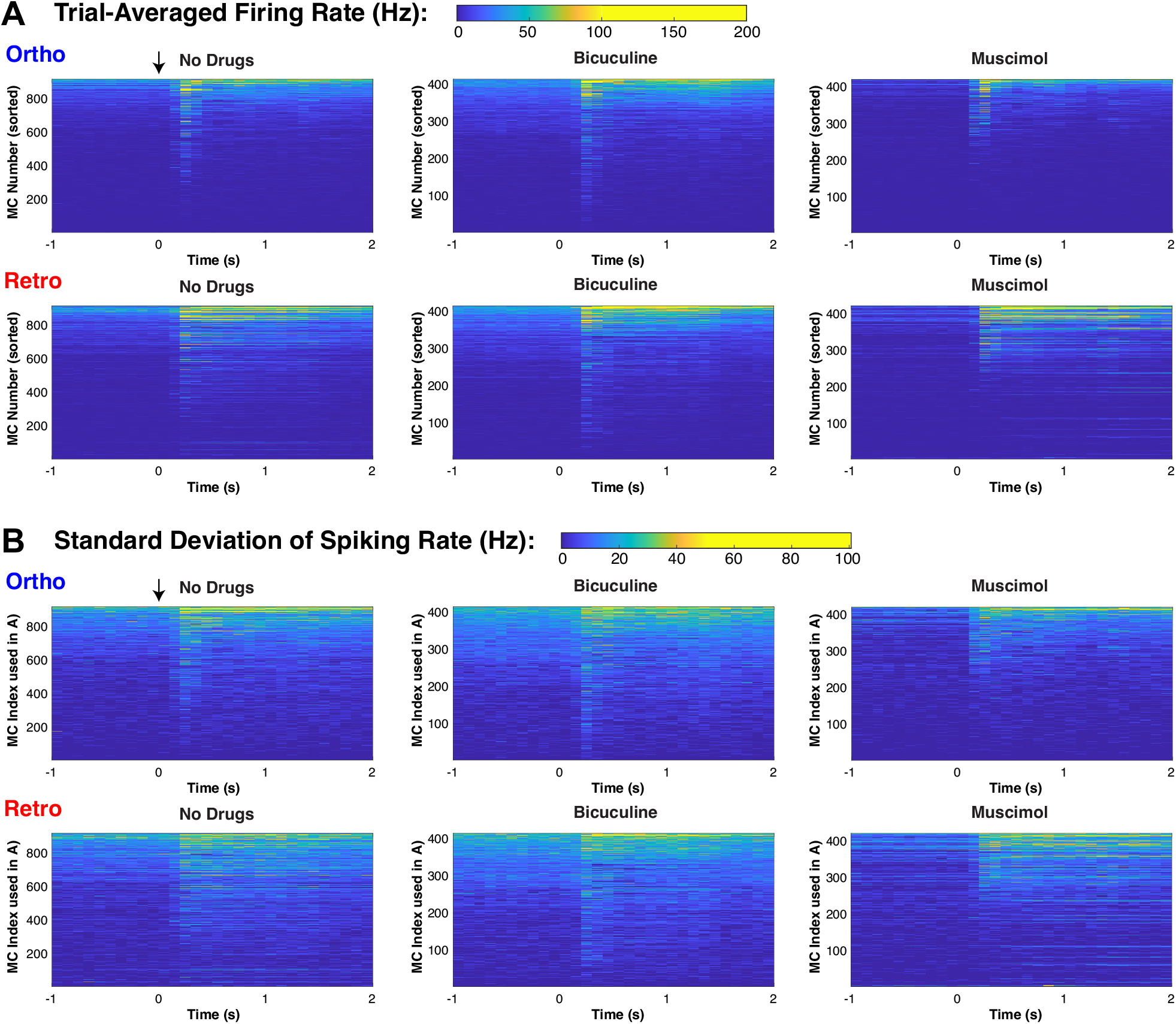
Detailed spiking rate statistics of pooled MC populations. **A**) Colormap showing trial-averaged spiking rate of individual MC (individual PSTH), each of the 3 columns is sorted by time-averaged firing rate (4 s total, 2 s in spontaneous and evoked, but only showing 1 s of spontaneous for brevity) of ortho firing rate, so using 3 ways to sort. **B**) Colormap showing trial variability of spiking rate (measured by standard deviation across trials) of individual MC, using the same corresponding index in top row (ortho firing rate). To see relative differences across MC, the color scales were effectively thresholded so that the larger values did not obscure the entire population. See Fig A3 for more on firing rate and trial-to-trial variability relationship.

### Drug effects on average population spiking

To better understand the dynamics of MC decoding accuracy, we consider how GABA_A_ (ant)agonists effect population activity. Although population activity is a coarse measure that neglects individual cell heterogeneity, it can still be insightful for determining average trends. A characteristic of an individual MC that might be indicative of encoding fidelity is the distance between the scaled trial-averaged firing rate of ortho and retro stimulation 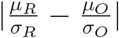(Fig 1B). We expect larger distances to coincide with better decoding accuracy than smaller distances, if all other factors are equal. Fig 3A shows moderate to strong correlations (Pearson’s and Spearman’s rank) between 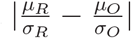 and decoding accuracies for all drug preparations. Although other factors affect decoding accuracy since the spread in the horizontal direction is large for a given decoding accuracy value and the correlations are not 100%. Nevertheless, this motivates considering the population-averaged spiking activity as a means to better understand the dynamics of decoding accuracy.

**Figure 3:**
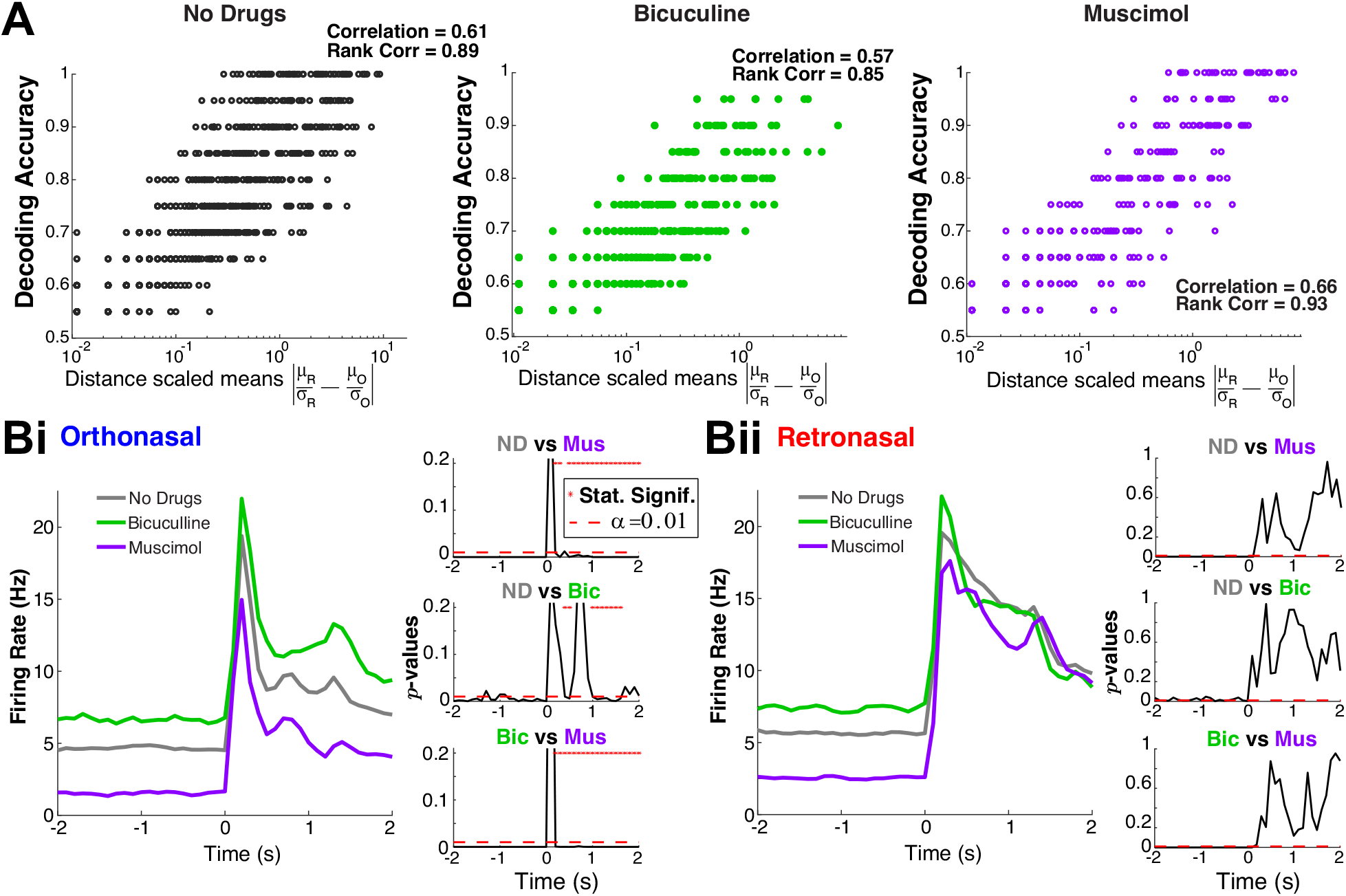
Population activity: retronasal response is more robust to changes in OB inhibition. **A**) Distance between scaled trial-averaged mean 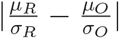 is correlated with decoding accuracy. The ‘Correlation’ is Pearson’s correlation coefficient, and ‘Rank Correlation’ is Spearmen’s. **B**) Drug effects on evoked population PSTH are different with statistically significant effects only with ortho stimulation (**Bi**) but not retro (**Bii)**. The right-panels with 3 subplots show the time series of *p* −values from two-sample *t* −test assuming unequal variances, comparing pairs of drug preparations.

The affects of the GABA_A_ (ant-)agonists on population firing rate depend on the odor modality; only ortho evoked firing rates show statistically significant differences in the mean (over MC population) firing rate (Fig 3B). As expected, Mus (increased inhibition) results in lower firing rates and Bic (less inhibition) results in higher firing rates compared to no drug, especially in the spontaneous state. However, the evoked firing rates are only significantly different (*α* = 0.01) across drug preparations with ortho stimulation, and not significantly different with retro stimulation. This holds for nearly all time points (see * in *p*−value plots of two sample *t*−test with unequal variance). This effect is clear in the right panels of Fig 3Bi and Bii. For completeness, the

Appendix shows the PSTH in Figs 1Aii and 3B for a longer timeframe (Fig A1A,B).

If we assume these population trends (Fig 3B) hold individually for MCs (i.e., drugs only effect spiking rate in ortho but not retro), then we might expect decoding accuracies to change in specific ways: if ortho firing rate is less than retro with no drug, then Bic should increase firing rate (shift histograms to the right) causing more overlap and lower decoding accuracy. For exposition purposes, these results are detailed in **Appendix** and Fig A4 where predictions on decoding changes based on individual MC’s relationship of ortho versus retro firing rate are partially verified.

### Drug effects on trial-to-trial variability

The trial-to-trial variability plays a key role in decoding accuracy differences. We analyze trial variance for each **MC** to test whether the population average is different across drug preparations. We find with Mus application, the average trial variability is smallest, followed by Bic, then no drug (i.e., **Mus** *<* **Bic** *<* **ND**), for both ortho and retro. The significance measured by *p*−values and effect size are shown in Tables 4 and 5 for ortho and retro trials, respectively, using the same three statistical tests as before. For ortho stimulation, Mus *<* ND is significant for two of the three tests (*α* = 0.01), while **Bic** *<* **ND** and Mus *<* Bic are only significant with Wilcoxon rank sum test. For retro stimulation, again only the Wilcoxon rank sum test shows this relationship is significant (all with *α* = 0.01 except for **Bic** *<* **ND** for retro where *p* = 0.038). Overall, the **Bic** *<* **ND** relationship is not as strong as the others, and the trend **Mus** *<* **Bic** *<* **ND** is stronger with ortho stimulation. Note that the effect size is considered ‘small’ in all cases, see Tables 4 and 5.

**Table 4:** Ortho stimulation only: significance (*p*−values) of average (over MC) trial variance differences between drugs with various statistical tests. Using same 3 tests as before; **ND**=‘no drug’; bottom table shows effect size.

| Relationship / $p$-value | $t$-test | Rank Sum | ANOVA |
| --- | --- | --- | --- |
| $\text{Mus} < \text{ND}$ | $1.8 \times 10^{-3}$ | $7.7 \times 10^{-8}$ | 0.021 |
| $\text{Mus} < \text{Bic}$ | 0.055 | $2.4 \times 10^{-13}$ | 0.053 |
| $\text{Bic} < \text{ND}$ | 0.82 | $5.0 \times 10^{-4}$ | 0.82 |

| Relationship / Effect Size | $t$-test | Rank Sum | ANOVA |
| --- | --- | --- | --- |
| $\text{Mus} < \text{ND}$ | 0.137 (small) | 0.147 (small) | 0.00402 (small) |
| $\text{Mus} < \text{Bic}$ | 0.134 (small) | 0.254 (medium) | 0.00450 (small) |
| $\text{Bic} < \text{ND}$ | 0.0137 (small) | 0.0955 (small) | $4.06 \times 10^{-5}$ (small) |

**Table 5:** Retro stimulation only: significance (*p*−values) of average (over MC) trial variance differences between drugs with various statistical tests. Using same 3 tests as before; **ND**=‘no drug’; bottom table shows effect size.

| Relationship / $p$-value | $t$-test | Rank Sum | ANOVA |
| --- | --- | --- | --- |
| Mus < ND | 0.039 | $1.9 \times 10^{-4}$ | 0.11 |
| Mus < Bic | 0.57 | $7.6 \times 10^{-7}$ | 0.57 |
| Bic < ND | 0.26 | 0.038 | 0.30 |

| Relationship / Effect Size | $t$-test | Rank Sum | ANOVA |
| --- | --- | --- | --- |
| Mus < ND | 0.0953 (small) | 0.102 (small) | 0.00193 (small) |
| Mus < Bic | 0.0399 (small) | 0.171 (small) | $3.98 \times 10^{-4}$ (small) |
| Bic < ND | 0.0610 (small) | 0.0569 (small) | $7.97 \times 10^{-4}$ (small) |

Within all of the 3 drug preparations, the average trial variances for ortho compared to retro are statistically indistinguishable, consistent with Craft et al. (2021) who did similar analyses but on only 1 rat instead of 8 rats.

### Model connects network dynamics and decoding accuracy

To reveal the neural network dynamics that explain our experimental data results, we use a simple firing rate model. Our rich data set with 3 total drug preparations enables a model framework that is highly constrained by data (Xiao et al., 2021; Ly et al., 2021; Barreiro et al., 2017). In addition, we incorporate known differences in temporal dynamics of ORN inputs to MCs with ortho versus retro stimulation (Sanganahalli et al., 2020; Gautam and Verhagen, 2012b; Furudono et al., 2013; Scott et al., 2007; Carey et al., 2009). Ortho stimulation results in fast increase and fast decrease of ORN inputs to MCs while retro input results in slower increase and slower decrease ORN inputs than ortho (Fig 4B). Fig 2c of Furudono et al. (2013) suggests that the amplitude of ORN inputs with ortho are larger than retro (Fig 4B). We develop a principled model that is structurally the same with both odor modalities but effectively have different dynamics stemming from different ORN inputs (Craft et al., 2021); note that paper had a different purpose with data only from intact circuit, i.e., it did not include drug manipulations. We decided not to use the biophysical model in Craft et al. (2021) because: i) individual MC spiking statistics are modulated by net inhibitory inputs, distinguishing interneurons is a complication more relevant for joint statistics of MC populations; ii) resolving trial-average spike statistics is more computationally expensive, making parameter tuning to capture *both* firing rate differences and average decoding accuracies across 3 drug preparations impractical; iii) calculating many decoding accuracies where each parameter set has an optimal threshold that has to be determined exhaustively (see Fig 5B,C below) is a necessary additional component here (not in Craft et al. (2021)) that is also computationally expensive.

**Figure 4:**
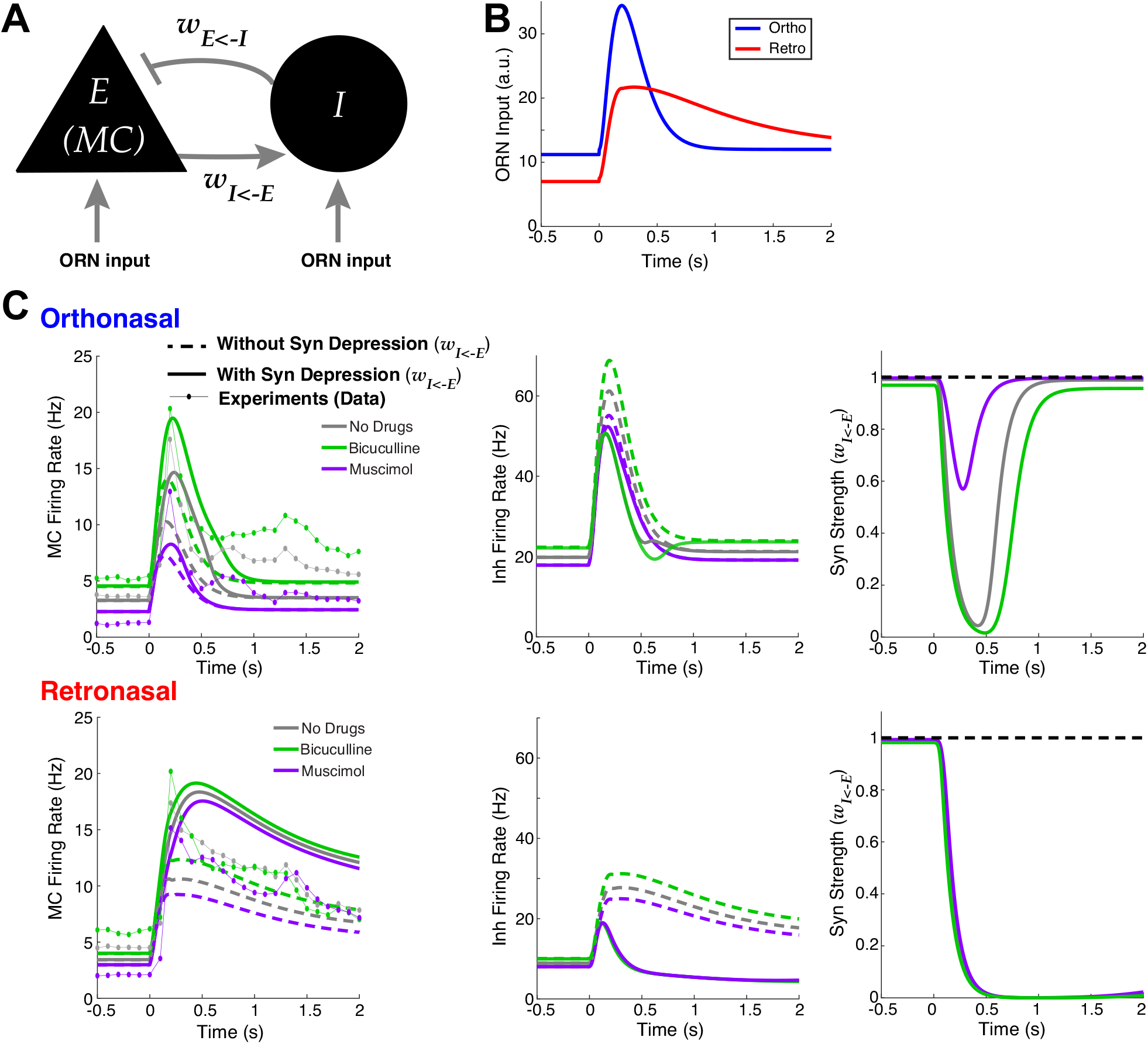
Firing rate model captures effects of drugs, with better alignment with data when including synaptic depression. **A**) Schematic of reciprocally coupled E-I pair. **B**) Fixed ORN input profiles for ortho and retro based on experiments (Sanganahalli et al., 2020; Gautam and Verhagen, 2012b; Furudono et al., 2013; Scott et al., 2007; Carey et al., 2009) (see main text). **C**) Wilson-Cowan firing rate model (Eqs (6)– (12)) output for ortho (top row) and retro (bottom row) stimulation, compared when synaptic depression is included (solid lines) or not (dashed lines). Left column: *E* cell (**MC**) output; center column: *I* cell output; right column: synaptic strengths (*W*_*IE*_). Data from experiments also shown for comparison (dot-lines) in left column.

**Figure 5:**
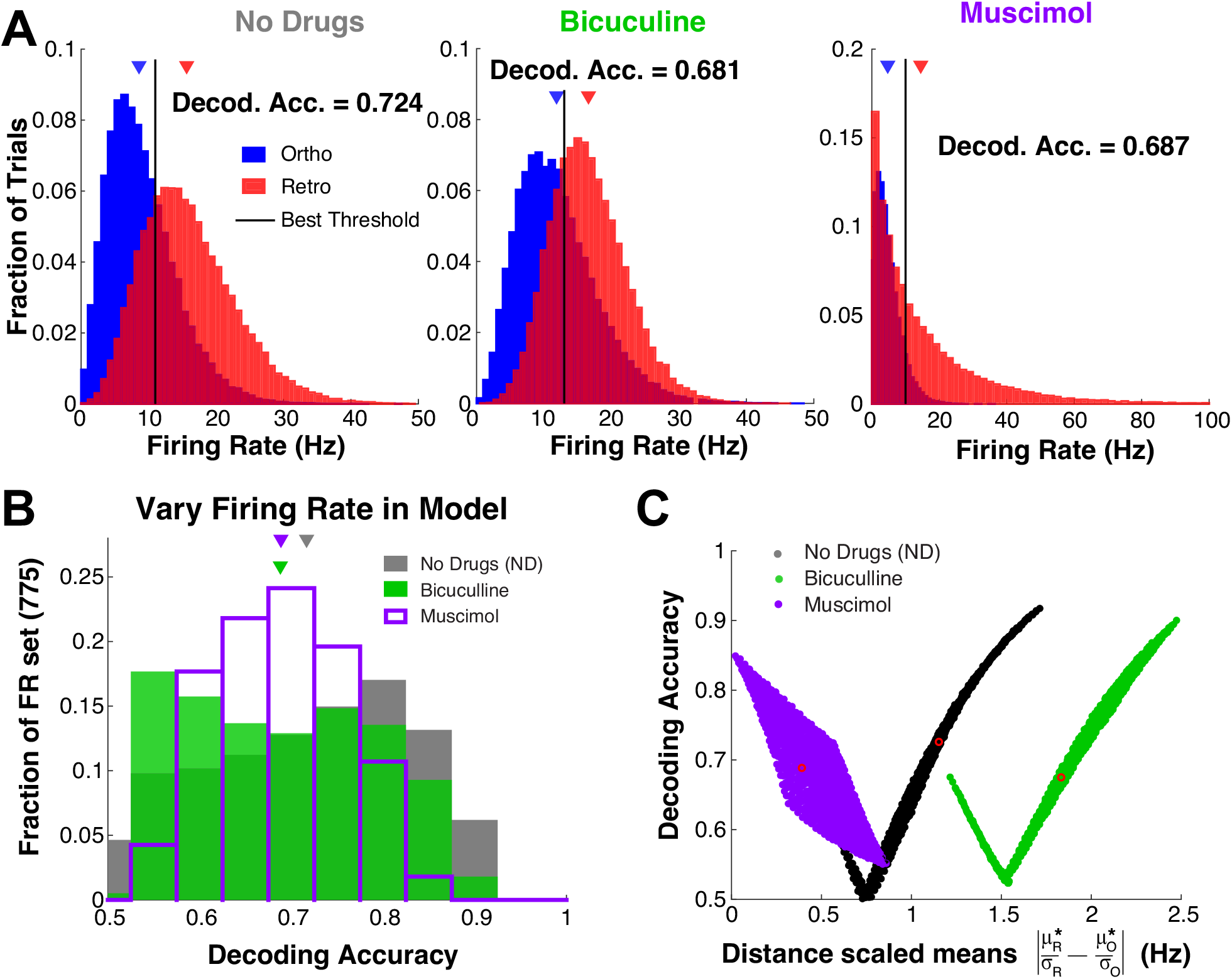
Model decoding accuracies with simulated trial variability. **A**): Histograms of simulated firing rate assuming negative binomial trial-to-trial variability with the mean set to: 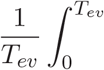 (see Eqs (6)–(12), and solid curves in left column of Fig 4 for *A*_*E*_(*t*)). **B)–C**) Systematically varying the mean to show that the specified *ρ* parameter (Table 1) in negative binomial distribution generally results in no drug having best decoding accuracy (see main text for details). **C**) Same model results in **B**) but showing relationship with distance between (scaled) means; red circles correspond to regimes in **A**).

**Figure 6:**
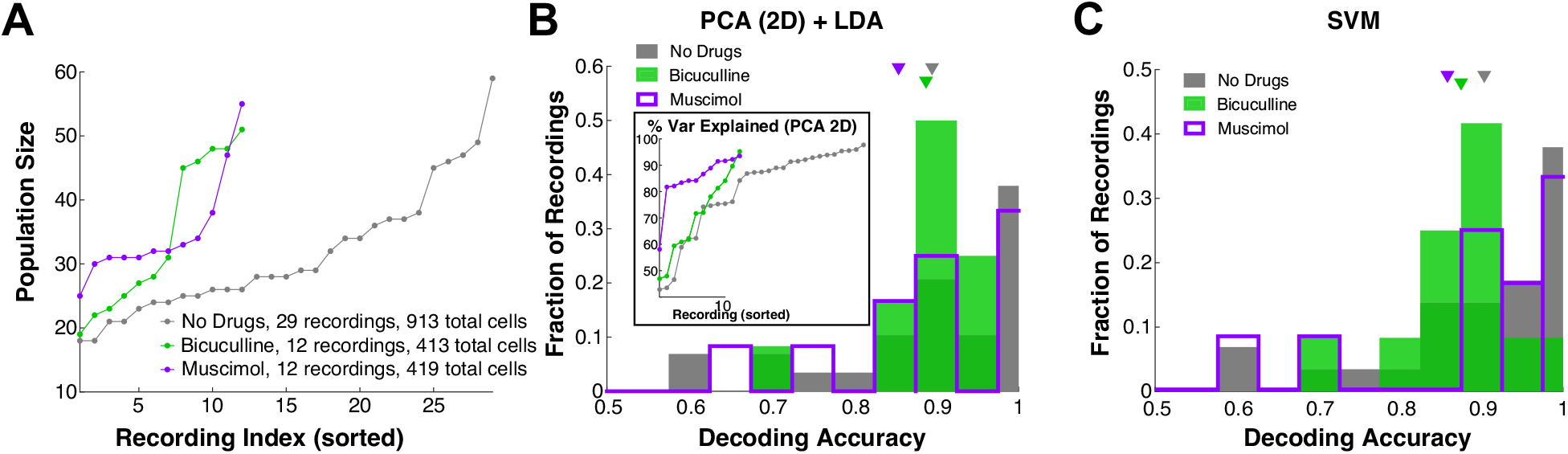
Population decoding of ortho versus retro is reliable and robust to altered inhibition. **A**) Each drug preparation has a different number of recordings and each recording has a different number of simultaneously recorded MCs. Showing recording number (sorted) and MC population size. **B**) Decoding accuracies (one for each simultaneous recording) for all drug preparations using a 2D PCA dimension reduction following by LDA. Inset shows variance explained by PCA (sorted) by recording. **C**) Decoding accuracies using full dimensional support vector machine (**SVM**) classification. Average decoding accuracies by both methods (**B)–C))** are statistically indistinguishable (minimum *p*−value for all relationships and all 3 statistical tests is: *p* ≥ 0.32 for PCA+LDA, *p* ≥ 0.13 for SVM).

We first set out to implement an OB model that accounts for the large changes in population PSTH with GABA_A_ (ant-)agonists with ortho stimulation <u>only</u>, and small (or no significant) changes in population PSTH with retro (Fig 3B). We use a reciprocally coupled 2 cell E-I network (Fig 4A), consistent with OB networks that have many such E-I pairs (granule and periglomerular cells are reciprocally connected to MCs with fast dendrodendritic synapses (Rall et al., 1966)). We model the drug effects Bic (Mus) by decreasing (increasing) the coupling strength from I to E cell by 25% of the baseline value (*w*_*EI*_ = 0.2 with no drugs, arbitrary units, see Eqs (6)–(7) and Table 1).

The data shows that affects of drugs altering inhibition are largely absent with slower longer-lasting retro stimulation (Fig 3B), thus finding a neural network principle that effectively removes the inhibitory cell input is imperative, and might be indicative of the core mechanisms. We hypothesize that short-term plasticity on the excitatory synapse (*E* → *I*) could be a factor in explaining the differences of population firing between ortho versus retro with GABA_A_ (ant-)agonists because of known temporal differences in ORN inputs. There is indeed evidence of plasticity in the OB circuit (Cang and Isaacson, 2003), specifically for short-term synaptic depression in the main OB (Wang et al., 2012) and from MC to granule cells (Dietz and Murthy, 2005). Since the relative differences in firing rate with ortho are already large without plasticity (1st column in Table 6), honing in on decreasing the firing rate differences in retro (3rd column in Table 6) would make the model more consistent with the data (Fig 3B). We use short-term synaptic depression as a means to nonlinearly change the retro firing rates to diminish the differences because retro has longer lasting stimuli. Although Dietz and Murthy (2005) showed that *E* → *I* synapses in OB have both facilitation and depression with facilitation appearing more prominent, we found that short-term facilitation alone made our model much less consistent with our data.

**Table 6:** Relative differences of firing rate model between no drug and inhibitory d rugs in both modalities. All /percent differences (top table) are relative to no drug: 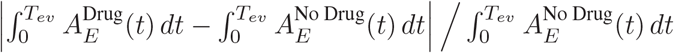 Bottom table shows average firing rate^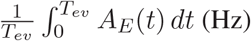^ i.e., the components of the relative differences.

| Relative Differences (%) | Ortho |  | Retro |  |
| --- | --- | --- | --- | --- |
|  | Without Depression | With Depression | Without Depression | With Depression |
| No Drug / Bicuculline | 37.87 | 38.16 | 16.08 | 6.04 |
| No Drug / Muscimol | 30.31 | 45.09 | 12.90 | 6.25 |

| $\frac{1}{T_{ev}} \int_0^{T_{ev}} A_E(t) dt$ | Ortho | | Retro | |
| --- | --- | --- | --- | --- |
|  | Without Depression | With Depression | Without Depression | With Depression |
| No Drug | 6.11 | 8.58 | 9.64 | 15.65 |
| Bicuculline | 8.43 | 11.86 | 11.19 | 16.59 |
| Muscimol | 4.26 | 4.71 | 8.40 | 14.67 |

Figure 4C shows comparisons of various model outputs (Eqs (6)–(12)). The model indeed follows the firing rate modulation with GABA_A_ (ant-)agonists with both odor modalities, whether or not synaptic depression is included. Specifically, ortho MC firing rates (top left in Fig 4) with different drug preparations are well-separated, while the retro MC firing rates are pretty close together (bottom left in Fig 4). The effects of synaptic depression significantly narrows the differences in retro population PSTH among all 3 drug applications, while with ortho the differences across drug preps remain large (see Table 6 for relative differences). Taken together, synaptic depression is crucial for capturing the data (Fig 3B). With retro stimuli and all 3 drug preparations, the *E* → *I* synaptic connections are depressed for longer so that the *I*-cell fires less, giving a boost to *E*-cell firing. In both modalities, there is overall higher *E* (MC) firing with short-term synaptic depression because of less inhibition (for all 3 drug preparations). Synaptic depression effectively diminishes the reciprocal loop so that the inhibitory drug effects are weakened. In principle Mus should have the opposite effect as Bic, but: the coupling strength *w*_*EI*_ has moderate variation: 0.15 (Bic), 0. 2 (ND), 0.25 (Mus). Also, we modeled short-term depression so that inhibition is almost <u>completely</u> turned off with Retro, resulting in MC being effectively decoupled from the *I* cell. Our model does not have realistic levels of synaptic depression, Cang and Isaacson (2003) report moderate levels of synaptic depression. If synaptic depression was limited in our model so that the minimum *w*_*IE*_ was 0.5, the ortho firing would be similar but the retro firing would strongly modulate with drug effects, which is inconsistent with our data. And anesthetized rats have less active granule cells (Cang and Isaacson, 2003) than awake, so perhaps other interneurons would have to be active for our model to be more consistent. The exact type of inhibitory neurons in the OB that would have to be involved and their corresponding dynamics with excitation are unknown to us.

With the neural dynamics captured in our model, we turn to connecting the model to decoding accuracy. The firing rate model does not explicitly have trial-to-trial variability, an important piece for decoding accuracy. A natural choice would be to assume Poisson statistics for the spike counts, but this single parameter distribution results in trial variance ordered by the firing rate: Mus*<*ND*<*Bic, which is <u>inconsistent</u> with our data and has perfect (100%) decoding accuracy with these firing rate models. Capturing the trial variability trends in our data seems to require tunable parameters that are not directly tied to the (mean) firing rate. We use a simple two parameter model that makes simulation easy and transparent: the negative binomial distribution. Note that there may be other models of trial variability that are more realistic relation to MCs, but that is unlikely to be as pragmatic for capturing firing rate <u>and</u> decoding accuracy trends. Specifically, for each drug (ND/Bic/Mus) and each input (ortho/retro), we fit a negative binomial distribution to the average decoding accuracy. The two distribution parameters were chosen so that: 1) the mean of distribution coincided with the model spiking rate 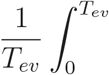 (solid curves in Fig 4C, left column), and 2) the decoding accuracy matches the averages in the experimental data.

The negative binomial distribution has distribution: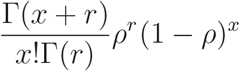with parameters: *ρ* ∈ (0, 1) and *r >* 0 that are not directly related to the physiology. We manually (by trial and error) determine the parameter *ρ* for each modality and drug preparation, which are all the same *ρ* except for retro with Bic and Mus (see Table 1), after which the *r* parameter is determined by:

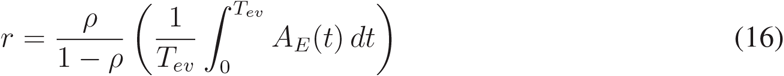

since the mean of the negative binomial is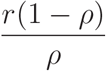We chose *ρ* for each regime so that the resulting decoding accuracy is close to the averages from the experiments: 0.724 (no drug), 0.680 (Bic), and 0.690 (Mus).

The results are summarized in Fig 5A. We simulated 50,000 trials and found the optimal threshold to get decoding accuracies that match the data.

We check whether our model results are consistent with the trial-to-trial variability of our data by calculating the variances of firing rate over trials, which is simply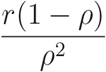 for the negative binomial distribution. The values listed in Table 7 show that trial variance is consistent with the data: Mus *<* ND *<* Bic, except for retro Mus firing that has the largest variance. Recall in our data that the trends for variance of firing rate was weakest with retro stimulation with larger *p*−values (Table 5).

**Table 7:** Simulated trial variance of the model: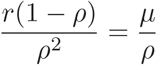.

|  | Muscimol | No Drug | Bicuculline |
| --- | --- | --- | --- |
| Ortho | 14.73 | 26.83 | 37.06 |
| Retro | 282.12 | 48.90 | 31.31 |

A test of our model is to determine whether the specified parameter values for *ρ* results in ND generally having better decoding accuracy than Bic and Mus. We start with the base model of spiking rate:

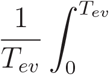 (solid curves in Fig 4C, left column) and perturb these values to get different means: 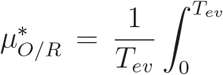 where *M*_*O*_ ∈ {−2, …, 4} Hz on 25 equally spaced points and *M*_*R*_ ∈ {−6, …, 9} Hz on 31 equally spaced points. Once a mean is specified, we use the same *ρ* (Table 1) and set *r*^∗^ to correspond to the mean *µ*^∗^:

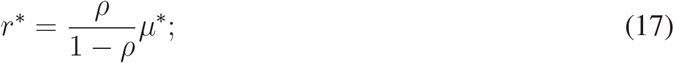

we simulate 50,000 trials to get a decoding accuracy value for each set of parameters. We get 775 decoding accuracy values corresponding to the different 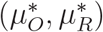 combinations for each drug preparation (Fig 5B). Importantly, our model does not soundly capture many details of realistic MC heterogeneities in our data (Fig 1C and Fig 5B have different shapes), but Figure 5B at least matches the result that ND on balance has better decoding accuracy than Bic and Mus. Finally, Figure 5C shows how decoding accuracy varies with scaled distance 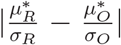The decoding accuracies can vary for a given drug preparation, but the ND and Bic have strong positive correlations (like the data, see Fig 3A), but Mus has strong negative correlations that is inconsistent with the data.

### Population decoding results

Thus far we have focused on decoding odor modality using the firing rate of individual MCs. While this approach avoids making assumptions about how higher level cortical regions might use the population as a whole (Rolls and Treves, 2011), population coding is known to be important for complicated tasks (Saxena and Cunningham, 2019). We apply standard population coding metrics to assess how well odor modality can be decoded from populations of MCs. We show that modality is well-encoded, but that the differences with GABA_A_ maniuplations are minor.

We applied two approaches to population decoding. First, we used principal component analysis **(PCA)** to project the population response onto a two-dimensional subspace that captures the most variance (first 2 PCs), and then applied linear discriminant analysis (**LDA**) to find an optimal decoder in that subspace (PCA+LDA). Second, we used a support vector machine (**SVM**), a common supervised learning algorithm to (nonlinearly) classify data. We use the first 2 PCs in PCA+LDA because it is a natural extension of 1D decoding with LDA that was the primary focus of prior sections. The SVM is at another extreme using all dimensions possible; together, these two provide a glimpse into the various accuracies of population decoding.

With both methods, we find odor modality is still encoded well above chance even when inhibition is altered (Fig 6B,C). Not only is the average decoding accuracy (across separate recordings) higher than before (Fig 1C) but there are no longer decoding accuracies near chance level (0.5). This is perhaps expected because these metrics (PCA+LDA, SVM) exploit the higher dimensional population activity to perform optimally in a binary classification problem when in reality the MCs encode other aspects of stimuli. Note that nearly 35% of recordings with no drug and Mus have perfect decoding accuracy (Fig 6B,C).

In contrast to the case of a single MC’s encoding (Fig 1C), we do not find any statistically significant differences between the mean decoding accuracy across drug preparations. In fact, the minimum *p*−value for all relationships and all 3 statistical tests is *p* ≥ 0.32 for PCA+LDA, *p* ≥ 0.13 for SVM. We expect population coding to perform better than individual cell coding, especially given the large dimensionality of population activity and that these metrics (PCA+LDA, SVM) were optimized for a binary classification task.

## Discussion

We have shown that MCs in the OB encode odor modality with our experimental data, and use an OB model inspired by many prior studies and constrained by our rich data to analyze plausible network components that support these dynamics. Our study uncovers further details for how ortho versus retro information is encoded to early cortical regions via the OB, which is consistent with reports that humans are able to discriminate whether food odors are delivered ortho or retronasally without being told which modality (Frasnelli et al., 2008). Imaging studies of the human mouth have shown that with food odors, ortho stimuli delivery versus retro are two distinct modalities, rather than 2 different stimulus routes to the same modality, independent of odor intensity (Bender et al., 2009). Based on *in vivo* rat data, we found significant differences in decoding accuracies (our proxy for encoding assuming an ideal observer) for classifying ortho and retronasal odors across different drug preparations. The intact (no drug) circuit had the best (average) decoding accuracy, suggesting that the OB might be structured to encode this important aspect of odor input.

Our model framework used prior results, data constraints (Xiao et al., 2021; Ly et al., 2021; Barreiro et al., 2017), and simplicity to account for the results in our data. Many prior studies (Sanganahalli et al., 2020; Gautam and Verhagen, 2012b; Furudono et al., 2013; Scott et al., 2007; Carey et al., 2009) showed that there are differences in the temporal dynamics of ORN response to ortho versus retro stimulation. We used this to investigate the OB circuit components that promote efficient coding of individual mitral cells with drug manipulations of inhibitory synapses. Further data analysis revealed that inhibitory drugs have a stronger effect on the population firing with ortho stimulation than retro. Using these insights, we constructed a standard firing rate model that captures the various drug effects for ortho versus retro stimulus, using only a pair of *E* − *I* cells. Including synaptic depression was crucial for capturing the different drug effects of firing rate. We did not explore varying the many parameters of the model to find a ‘best fit’ to data, rather we wanted a simple yet informative model that demonstrates a plausible mechanism. The model PSTH fits to the data are imperfect and qualitative (Fig 4C): the spontaneous firing of ortho and retro in the model are different because of differences in the ORN input strengths, the ortho model firing decays faster than the data. In fact for retro it looks like the model fits the data better without synaptic depression (although the differences with GABA_A_ manipulations are large and inconsistent with the data). In theory the model could be better fit to the data PSTH, for example if the ORN inputs (Fig 4B) are held fixed and an optimization routine is performed on a set of parameters, but then the decoding accuracy would have to be accounted for (Fig 5A), perhaps in the objective. We thus opted to manually determine the model parameters. Lastly, we simulated trial-to-trial variability by assuming a negative binomial distribution with mean defined by the firing rate model output. The resulting model captured both decoding accuracy trends (ND is better) and some aspects of firing rate trial variability.

There have been prior studies that implemented an OB circuit model with varying levels of realism, but none that we are aware of that uses an OB model to account for coding of odor modality and firing rate dynamics. We previously developed a realistic biophysical OB network model (Craft et al., 2021) and used it to account for different neural network dynamics (trial-averaged) with ortho/retro inputs, but did not focus on decoding accuracy for a given trial (which is more computationally taxing). In our prior work (Craft et al., 2021), the PSTH for ortho versus retro are different than here (Figs 1Aii, 3B) because we previously used only 1 rat versus 8 rats here. There are biophysical models of particular OB cells (Viertel and Borisyuk, 2019; Li and Cleland, 2013) and of the OB network (Bathellier et al., 2006; Li and Cleland, 2017), and other have used models (firing rate or Bayesian) to investigate coding of mixed odors (Grabska-Barwińska et al., 2017) and learning new odors (and their concentration levels) (Hiratani and Latham, 2020), but none that we know of that focus on coding of odor modality. Note that muscimol increases tonic E cell inhibition independent of I cell firing; our simple model does not disentangle these two components.

A limitation of this study is the use of one odor (EB, a food odor) with one concentration. We collected data from a non-food odor (1-Hexanol), but since typically only food odors experienced retronasally (Small et al., 2005), we did not include 1-Hexanol in our study. It is well-known that non-food odors delivered retronasally are not perceived as well as retro food odors by humans (Hannum et al., 2021; Bender et al., 2009; Frasnelli et al., 2008). Other food odors have different physicochemical properties that impact differences in ortho versus retro responses (Scott et al., 2007), so the generalization of our results merits further investigation. Firing rates vary with odor concentration (Tan et al., 2010) and is thus a confounding factor in distinguishing ortho vs retro from firing rates alone. Whether higher brain regions rely on other signals to account for concentration or different patterns of activation in OB to reliably estimate ortho/retro is important but beyond the scope of this study. Our data was collected from anesthetized rats with forced air to model ortho/retro (Gautam and Verhagen, 2012b,a; Gautam et al., 2014) which has clear advantages: control and a fair decoding ‘task’ so that ortho and retro are mechanically similar with same stimulus duration. In addition to the intact circuit, our data also had both increased and decreased GABA_A_ synaptic inhibition with direct manipulations in the OB. Via pooling data from many rats, we had a large number MCs. Another limitation is that the rats were not awake and were not eating. We are currently unaware of any data with awake rodents that record MC spiking activity for the purposes of comparing ortho and retro (food) odors during feeding. There are freely available MC spike data with awake rodents (mice in Bolding and Franks (2018, 2021)) but not during feeding and not with direct manipulation of OB inhibitory synapses.

Since the temporal dynamics of odors can be crucial in OB processing, we systematically varied the evoked time window and even considered removing the first 300 ms of the evoked state (see Fig A2). We found that the population-average (and standard deviation) decoding accuracy varied slightly (Fig A2B,D), and indeed with the first part of the evoked state removed the average decoding accuracy increased. In the context of our study, the temporal patterning differences between ortho and retro have minor effects on decoding. In awake, intact animals, the timing of the breath cycle determines the timing of ortho and retro stimulation (Shusterman et al., 2011). In principle, brain areas downstream from OB could use a breath-timing-based decoding scheme to entirely distinguish ortho from retro. However, our experiments show that timing is not the only mechanism that impacts ortho versus retro coding. In our experiments, the breath of the anesthetized rats did not pass through the nose at all; it was redirected through a tracheotomy tube. Thus, the timing of ortho and retro signals in our experiments was decoupled from breath timing and we kept the stimulation parameters (concentration and flow rate) the same, changing only the direction of airflow. Yet, we found differences in ortho and retro response in OB. Also, when instructed not to breath and actively sniff, humans can correctly report ortho versus retro odor modality with food odor without being told the modality (Frasnelli et al., 2008). Thus, we conclude that the OB encodes ortho and retro stimuli not just with the timing of the breath, but also with other mechanisms. Nonetheless, it is likely that intact, awake animals would employ both timing (inhale vs exhale) and the mechanisms we identify here. Disentangling the relative importance of these different possible contributions to ortho versus retro coding will require further experiments. Our results predominately center on population averages of individual MC decoding accuracy because it is a logical first step (Rolls and Treves, 2011). Given the importance of population coding for many complicated tasks (Saxena and Cunningham, 2019), we also considered 2 standard population decoding accuracies measures: PCA+LDA and SVM and found in both cases that the intact circuit was not statistically significant better (on average). However, decoding accuracy was again much higher than chance levels and better compared to average individual MC decoding accuracy. Developing a heterogeneous OB model that accounts for the population coding results and dynamics is a potential next step but beyond the scope of this current study.

## Author Contributions

MC programmed and implemented models in software, including the data analysis. MC and CL developed the computational models. MC, AKB, WLS, CL conceptualized and developed the project. SHG and WLS designed the experiments and collected the data. MC and CL drafted the original manuscript, including visualizations. MC, AKB, SHG, WLS, CL edited the manuscript. CL supervised the project.

## Appendix

The main text focuses on the 1 s duration of the odor, which is longer than most breath cycles. However, there are dynamics beyond the duration of the odor that may be insightful and of interest, so the population PSTHs extended to 5 s after odor onset are shown in Figures A1A,B. By visual inspection, there appears to be an increase in the firing rate after the odor is off, e.g., ortho with Bic and retro with Mus. Whether this change is statistical significant, and the underlying mechanisms to explain this are fascinating but beyond the scope of this current study. For the interested readers, we report the fraction of ‘Off-Response’ cells that have higher firing rates when the odor is removed than with odor, a well-known phenomena dating back many decades (Meredith, 1992). We report the fraction of the MC population the has a higher firing rate (time and trial-averaged) in 1 ≤ *t* ≤ 2 s than in 0 ≤ *t <* 1 s in Table A1.

**Table A1:** Fraction of ‘Off-Response’ cells: cells that have a higher firing rate (time and trial-averaged) in the immediate time (1 s) after odor is removed than during the 1 s odor stimulation.

| Fraction of Off-Response MCs | ND | Bic | Mus |
| --- | --- | --- | --- |
| Ortho | 30.12% | 36.08% | 27.45% |
| Retro | 27.05% | 18.64% | 42.00% |

**Table A2:** Correlation between trial-mean and trial-standard deviation (top) and variance (bottom) of spiking rate. There is a strong positive relationship that is more prominent with retro, see Figure A3. For further discussion on the modulation of evoked spiking variability in the OB, see Ly et al. (2021).

| Std. | Ortho |  | Retro |  |
| --- | --- | --- | --- | --- |
|  | Pearson’s | Spearman’s Rank | Pearson’s | Spearman’s Rank |
| **No Drug** | 0.932 | 0.986 | 0.935 | 0.995 |
| **Bicuculline** | 0.930 | 0.989 | 0.937 | 0.988 |
| **Muscimol** | 0.907 | 0.989 | 0.920 | 0.996 |
| Var. | Ortho |  | Retro |  |
|  | Pearson’s | Spearman’s Rank | Pearson’s | Spearman’s Rank |
| **No Drug** | 0.841 | 0.986 | 0.908 | 0.995 |
| **Bicuculline** | 0.835 | 0.989 | 0.963 | 0.988 |
| **Muscimol** | 0.852 | 0.989 | 0.968 | 0.996 |

**Figure A1:**
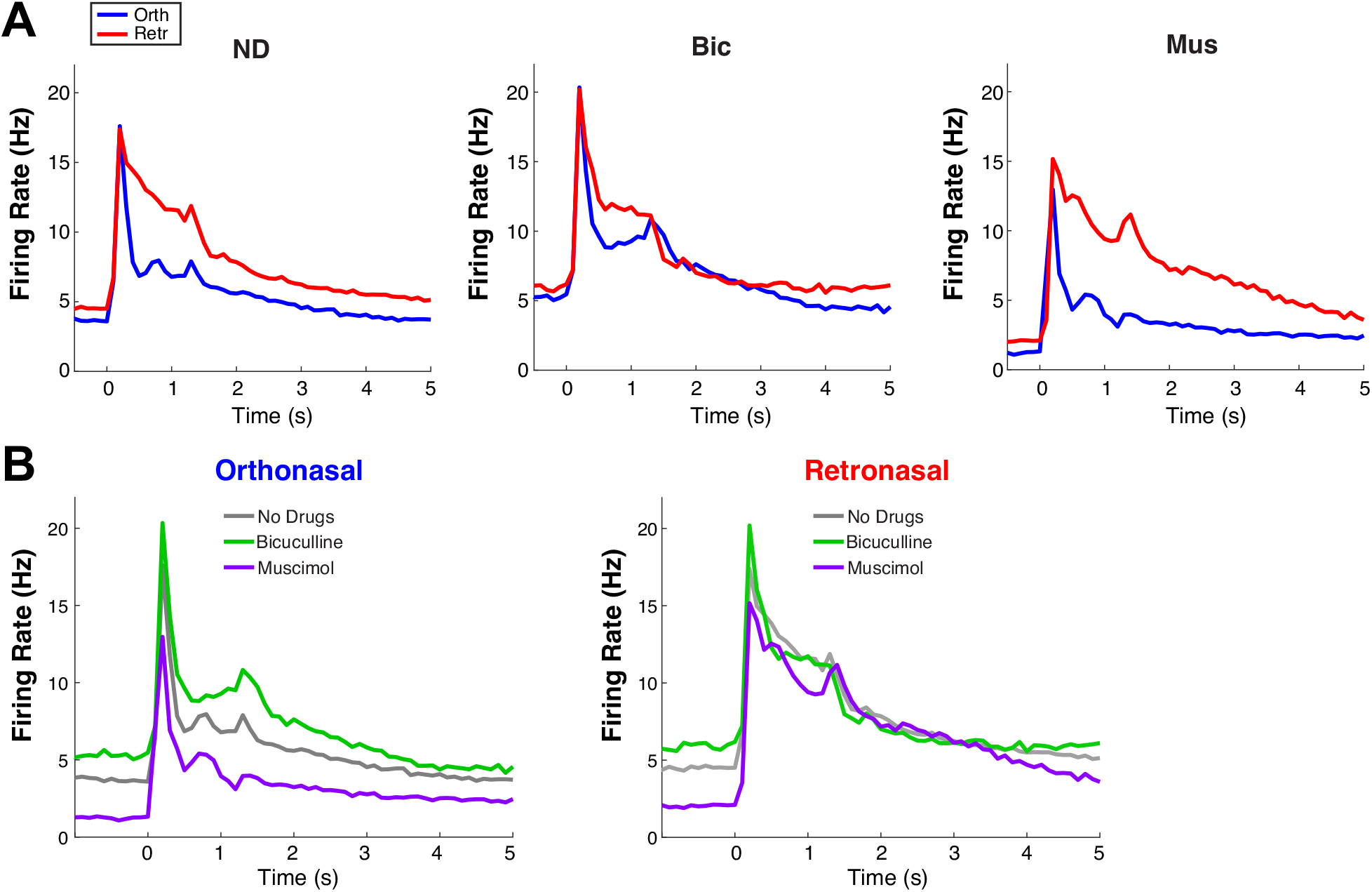
Supplemental data analysis: extended time of PSTHs in main text. Extended timeframe for Figure 1Aii (**A**) and Figure 3B (**B**).

### Decoding accuracy dynamics from drug effects on population-averaged firing rates

If we assume the population trends in Fig 3B) hold individually for MCs (drugs only effect spiking rate with ortho but not retro), then we might expect decoding accuracies to change in specific ways, as illustrated in Fig A4**Ai**,**Aii**. If ortho spiking is less than retro with no drug (*µ*_*O*_ *< µ*_*R*_, see Fig A4**Ai**), Bic should increase spiking (shift gray curve to the right, green curve) causing more overlap and lower decoding accuracy, while Mus should decrease spiking (shift gray curve left, purple curve) leading to less overlap and higher decoding accuracy. Similar arguments can be made for when retro spiking is less than ortho spiking (*µ*_*R*_ *< µ*_*O*_, Fig A4Aii). Thus, the predictions from observations of the population firing rates (Fig 2**B**) are:

1. When *µ*_*O*_ *< µ*_*R*_:
  a. Bicuculline = less accuracy*
  b. Muscimol = more accuracy
2. When *µ*_*R*_ *< µ*_*O*_:
  a. Muscimol = less accuracy*
  b. Bicuculline = more accuracy

The results of these predictions are shown in Fig A4**B**,**C**. Predictions 1a and 2a (Fig A4**B**) denoted by asterisk are statistically significant in difference (*t*−test (*p*_*ND, Bic*_ = 5.3×10^−16^, *p* _*ND,Mus*_ = 1.8 × 10^−6^), Wilcoxon rank sum test (*p* _*ND,Bic*_ = 3.1 × 10^−14^, *p* _*ND,Mus*_ = 6.9 × 10^−7^), and one-way ANOVA (*p*_*ND,Bic*_ = 1.1 × 10^−15^, *p*_*ND,Mus*_ = 5.6 × 10^−6^)). The other two predictions (1b and 2b, Fig A4C) do not precisely hold; interestingly, it is because no drug preparation retains a higher average decoding accuracy than with the two drug applications. Two caveats: in Fig A4**Ci**, with Mus the average decoding accuracy is better when *µ*_*O*_ *< µ*_*R*_ (purple filled) compared to all cells *µ*_*O*_ *< µ*_*R*_ & *µ*_*O*_ ≥ *µ*_*R*_ (dark purple outline) (*t*−test (*p* _*Mus*−*all,Mus*_ = 1.2 × 10^−8^), Wilcoxon rank sum test (*p*_*Mus*−*all,Mus*_ = 9.4 × 10^−6^), and one-way ANOVA (*p*_*Mus*−*all,Mus*_ = 3.0 × 10^−6^)); in Fig A4Cii, the difference between Bic and no drug are <u>no longer</u> statistically significant (*t*−test (*p*_*ND,Bic*_ = 0.22), Wilcoxon rank sum test (*p*_*ND,Bic*_ = 0.58), and one-way ANOVA (*p*_*ND,Bic*_ = 0.26)), so that with *µ*_*R*_ *< µ*_*O*_, cells with Bic and no drug have indistinguishable averages. This all shows that the observation that ortho stim is more effected with GABA_A_ manipulation than retro has consequences for decoding accuracy modulation, even if the predictions are imperfect.

**Figure A2:**
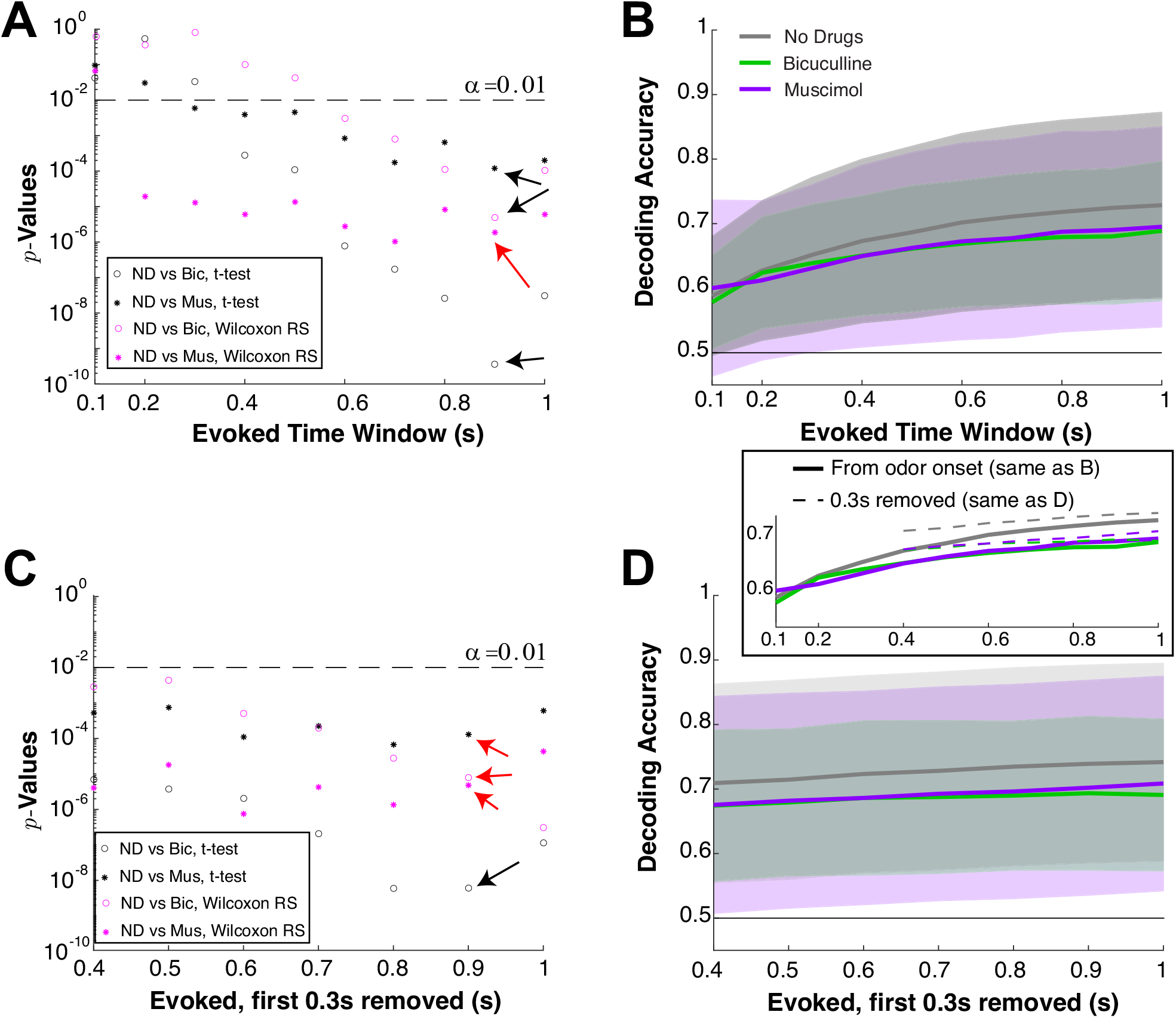
How we choose the time window; decoding accuracy as time window varies. Varying time windows on evoked firing rate to select for data analysis (**A, C**) and decoding accuracy. **A**) Vertical axes show *p*−values (log-scale) from comparisons of average decoding accuracies with different drug preparations using two different statistical tests: two sample *t*−test assuming unequal variance (black) and Wilcoxon rank sum test (magenta). Open circles compares no drug to Bic, stars compare no drug to Mus (see legend). **B**) Showing the population average decoding accuracy and ± one standard deviation in shaded region. Longer time windows generally lead to larger average decoding accuracy. **C), D)** are similar to **A), B**) except the first 300 ms of the evoked state are removed to test whether decoding accuracies increase are significant; this was based on the population PSTH being similar for ortho and retro immediately after stimulus onset (Fig 1Aii). Decoding accuracies are significantly different (*α* = 0.01, denoted by dashed black line) for a large range of time windows, but we chose 0.9 s for evoked because this value resulted in the smallest *p*−values generally, the exception being denoted by the red arrows where other time bins have *slightly* smaller *p*−values. We decided to keep the entire time, not excluding the first 300 ms, for simplicity and because the *p*−values are similar. All data analysis is with EB odor input.

**Figure A3:**
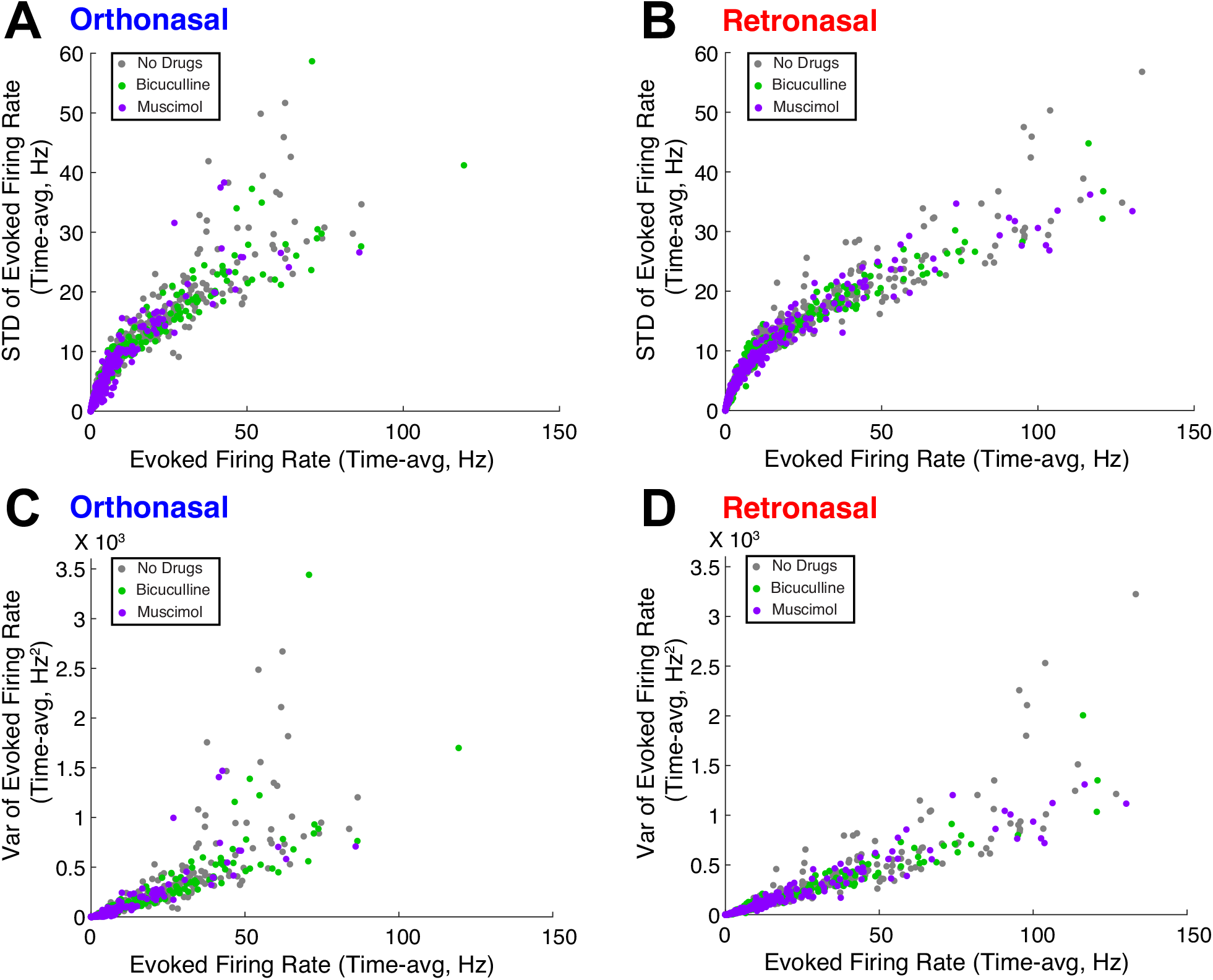
Trial-averaged firing rate is positively correlated with trial-to-trial variability. For each MC in a given modality and drug preparation, showing the trial-averaged firing rate versus the standard deviation (**A–B**) or variance (**C–D**) of firing rate (across trials). **A**) With orthonosal stimulation; **B**) with retronasal stimulation. **C)** and **D**) are similar except showing the variance of evoked firing rate. We plot the time average to resolve the many instances with no spiking so that each MC has a single number for the firing rate and a single number for the variance.

**Figure A4:**
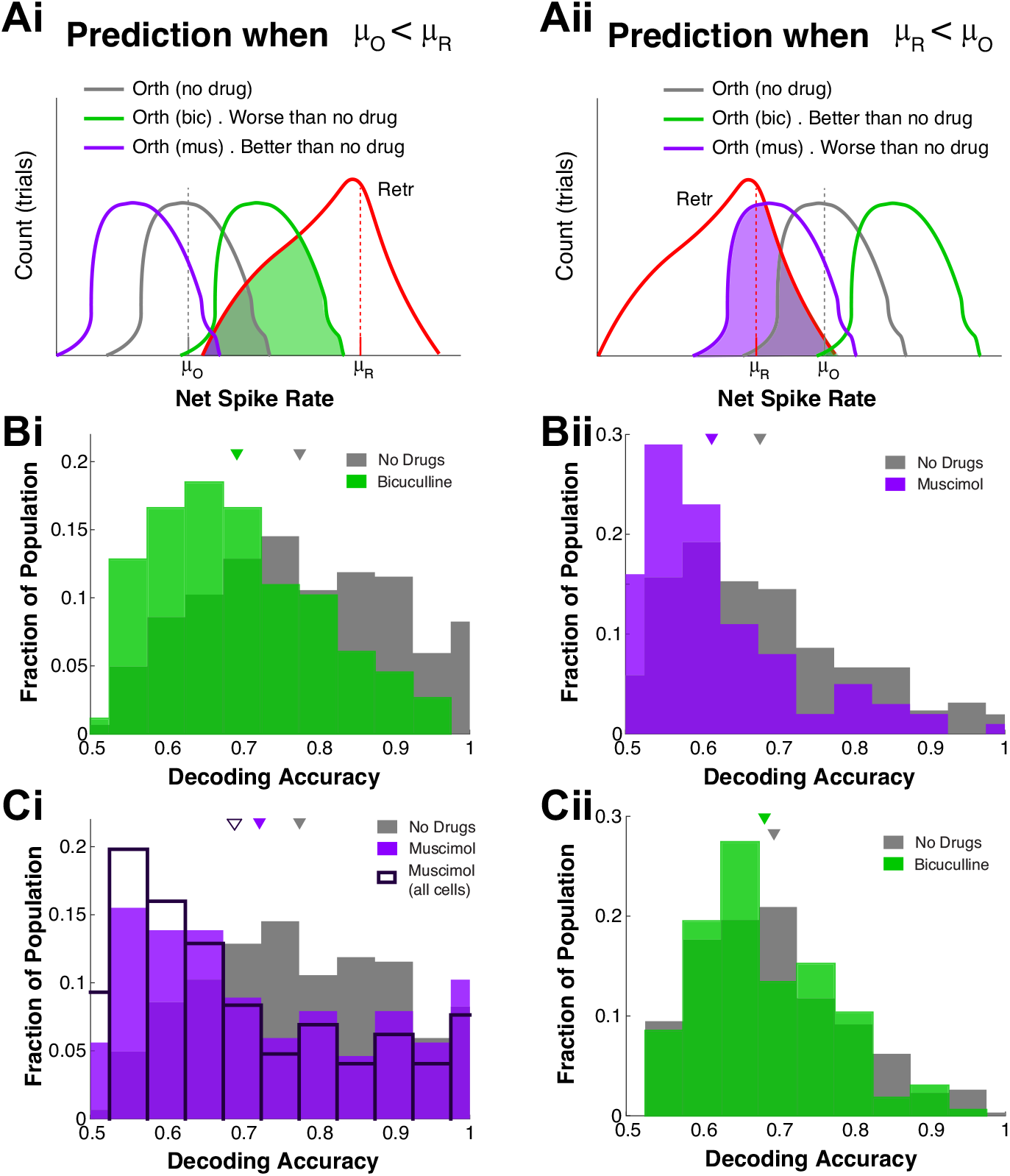
Decoding accuracy dynamics from drug effects on population-averaged firing rates. Left column: cells where *µ*_*O*_ *< µ*_*R*_; Right column: cells where *µ*_*R*_ *< µ*_*O*_. **Ai–Aii**) predicted decoding accuracy for different drug preparations. **Bi**) Histogram of decoding accuracies for no drug and Bic when *µ*_*O*_ *< µ*_*R*_, and no drug with muscimol when *µ*_*R*_ *< µ*_*O*_ (**Bii**); triangle is mean decoding accuracy over all such MCs. Differences in middle row (**Bi**,**Bii**) are statistically significant. **Ci**) Histogram of decoding accuracies for no drug and muscimol when *µ*_*O*_ *< µ*_*R*_, also showing the histogram for muscimol decoding accuracy for all cells (i.e., not only those where *µ*_*O*_ *< µ*_*R*_) in dark purple outline. Although no drug is better than muscimol, muscimol with cells *µ*_*O*_ *< µ*_*R*_ have better decoding than all cells with muscimol. **Cii**) No drug and Bic when *µ*_*R*_ *< µ*_*O*_ do not have statistically significant differences in mean. See text for *p*−values.

